# Measurement of tau protein and Aβ amyloid plaques in postmortem human brains of Down syndrome and Alzheimer’s disease by using [^125^I]IPPI and [^125^I]IBETA autoradiography

**DOI:** 10.64898/2026.01.17.700075

**Authors:** Agnes P. Biju, Fariha Karim, Deanna M. Schafer, Stephanie A. Sison, Christopher Liang, Elizabeth Head, Jogeshwar Mukherjee

## Abstract

The accumulation of tau tangles and Aβ plaques are prominent neuropathologies that characterize Alzheimer’s disease (AD) and Down Syndrome (DS). Continuous developments of PET tracers as biomarkers can be supported by autoradiography to validate effectiveness and accuracy of binding properties that elucidate the pathophysiology of DSAD and AD. This in vitro comparative study evaluates [^125^I]IPPI binding to tau and [^125^I]IBETA binding to Aβ plaques in the frontal cortex (FCX) and temporal cortex (TCX) of postmortem human brain slices of AD (n=5), DSAD (n=5), and cognitively normal (CN) (n=5) cases. With anti-tau and anti-Aβ immunostains confirming the presence of tau and Aβ plaques, [^125^I]IPPI and [^125^I]IBETA binding in autoradiographic images were significantly higher in DSAD and AD gray matter (GM) compared to CN. When comparing DSAD with AD, FCX and TCX GM binding was similar throughout DSAD and AD except in FCX GM where there was 48% more [^125^I]IPPI binding in DSAD than AD. In vitro drug inhibition studies revealed that [^125^I]IPPI binding was significantly inhibited with increasing harmine concentrations (IC_50_=115±40 nM) in DSAD FCX and TCX but KuFal194 minimally inhibited [^125^I]IPPI binding in the same cases. The GM/white matter ratios for DSAD ([^125^I]IPPI=4.1, [^125^I]IBETA=2.9) and AD ([^125^I]IPPI=4.2, [^125^I]IBETA=2.6) were significantly greater than CN ([^125^I]IPPI=1.3, [^125^I]IBETA=1.2). A positive correlation between [^125^I]IPPI and [^125^I]IBETA binding suggests a synergistic relationship between tau and Aβ plaque in DSAD and AD pathology. This study demonstrates that [^125^I]IPPI and [^125^I]IBETA may serve as novel radiotracers in both DSAD and AD to continue diagnostic investigations in vivo.

## 1. Introduction

Cholinergic terminal density decline, basal forebrain atrophy, and significant cerebral amyloid angiopathy are unique in people with Down’s syndrome (DS), with nearly all by the age of 40 years developing Alzheimer’s disease (AD) pathology (Rafii et al. 2025). The link of DS to AD is associated with the gene products resulting from an extra copy of chromosome 21 in DS, which can affect human physiology in multiple ways including dementia. There are differences between DS and AD, such as people with DS having greater tau concentrations than autosomal-dominant AD in the temporal lobe (Rafii et al. 2025). Because of other comorbidities (neurologic and psychiatric) accompanying DS, emerging molecular neuroimaging may be able to assist in improved diagnosis of DS-associated AD (DSAD) (Carmona-Iragui et al. 2019; Neale et al. 2017). The onset of AD in DS can depend on the individual’s resistance to AD pathology and resilience to pathological effects, complicating diagnosis strategies (Schworer et al. 2025) . Greater resilience to AD pathological processes influenced by lifestyle factors can delay the clinical symptoms of AD with cognitive functions intact that would otherwise be diagnosed as DSAD.

Accumulation of Aβ plaques occur in AD and has been successfully used as a biomarker (plasma, CSF, and neuroimaging) for disease progression (Zhang et al. 2023). Initial accumulation and brain regional distribution of Aβ plaques differ between AD and DS. Unlike in AD, accumulation of Aβ plaques in DS occurs in the striatum followed by frontal regions and lastly in the medio-temporal region (Annus et al. 2016; Abrahamson et al. 2019). Accumulation of Aβ plaques occurs in all adults with DS by the age of 40 years and can be noted as early as 35 years (Sokol et al. 2024). Striatal Aβ plaque accumulation is seldom seen in sporadic AD, but is observed in autosomal dominant AD, and neither has it been observed in transgenic mice models of such as 5xFAD which express abundant Aβ plaques at a young age. On the other hand, significant amounts of Aβ plaques are observed in the thalamus of 5xFAD mice using [^125^I]IBETA (Nguyen et al. 2022) and [^18^F]flotaza (Sandhu et al. 2024). Immunostaining suggests that thalamic Aβ plaques in mice are more diffuse compared to cortical Aβ plaques (Sandhu et al. 2024). Striatal Aβ plaques in DS appear prior to cortical accumulation and have been suggested to be primarily diffuse plaques (McLachlan et al. 2025). Significant levels of Aβ protofibrils and plaques in the inferior frontal gyrus and medial temporal gyrus in both AD and DSAD have been reported (Johannesson et al. 2021). Autoradiographic binding studies in postmortem AD brain slices with [^125^I]IPC-Lecanemab, which has a high affinity for protofibrils suggested presence of Aβ protofibrils (Liang et al. 2024). It is predicted that Lecanemab immunotherapy may be beneficial for people with DSAD, since they show Aβ plaque accumulation at earlier ages compared to AD (Hof et al. 1995). Age versus Aβ plaque load in adults with DSAD may be useful information in order to evaluate outcomes of therapeutic treatment protocols (Teller et al. 1996). Given that adults with DSAD have a significant amount of protofibrils and plaque, Lecanemab treatment may be beneficial when provided at younger ages.

Unlike Aβ plaques, regional brain tau accumulation in DS is similar to AD spreading from the hippocampus and entorhinal cortex to temporal cortex (TCX) and neocortex (Rafi, 2019). Since an interplay between Aβ plaques and neurofibrillary tangles (NFT) has been suggested, differences and similarities in the regional distribution of the two proteinopathies in DS and AD may provide additional biochemical information (Granholm and Hamlett 2024). Although early striatal Aβ plaque binding is associated with DS, no tau imaging agents have shown any significant binding in the striatum, either in DSAD or AD. This may raise the question of Aβ plaques being a progenitor for the formation of NFT at least in the striatum or they may be two independent molecular pathways (Liang et al. 2024; Perez et al. 2019).

Recent positron emission tomography (PET) studies using Aβ plaque imaging agents such as [^11^C]PIB (Lao et al. 2016) and [^18^F]florbetapir (Sabbagh et al. 2015) and tau imaging agents such as [^18^F]flortaucipir (Zammit et al. 2024) and [^18^F]MK-6240 (Hendrix et al. 2021) have provided insights into these differences between AD and DSAD. As mentioned previously, differences include the binding of Aβ plaque imaging agents in the striatum in DSAD. Postmortem studies of adults with DSAD with dementia show advanced NFT in the frontal cortex (FCX) but not in the striatal regions while Aβ plaque loads were similar in the two regions (Perez et al. 2019).

Recent cryo-EM structures of Aβ amyloid fibrils and tau filaments from DS show structures identical to those found in AD (Fernandez et al. 2024). Our previous molecular modeling and radioligand studies in human AD postmortem brains have confirmed the binding of [^125^I]IPPI to tau and [^125^I]IBETA to Aβ plaques (Mukherjee et al. 2021; Mondal et al. 2023). Due to the similarity between AD and DSAD Aβ plaques and tau filaments, [^125^I]IPPI and [^125^I]IBETA comparative binding in the FCX and TCX may provide useful information on the similarities and differences between these two pathologies. Such comparative studies in postmortem brain samples of AD and DSAD with the same radioligands are not available. Thus, we report autoradiographic and immunostaining studies of [^125^I]IPPI and [^125^I]IBETA in DSAD, AD and cognitively normal (CN) cases to compare their binding and expand upon the pathophysiological attributes of Aβ plaques and tau accumulation in FCX and TCX. We expect greater [^125^I]IPPI and [^125^I]IBETA binding in DSAD especially within FCX regions where AD pathology predominately starts.

## 2. Materials and Methods

### 2.1. General Methods

Iodine-125 labeled [^125^I]IBETA (Mondal et al. 2023) and iodine-125 labeled [^125^I]IPPI (Mukherjee et al. 2021) were prepared as reported previously. Various drugs, including harmine (Adooq Bioscience, Irvine, CA), MK-6240 and 1,3-bis(4-cyanophenyl)urea (1ClickChemistry, Inc, Tinton Falls, New Jersey, USA), KuFal194 (AABlocks LLC, San Diego, CA, USA) were purchased commercially. Capintec CRC-15R dose calibrator and Capintec Caprac-R well-counter were used for radioactivity measurements. Thin layer chromatography of radioligands were scanned on an AR-2000 imaging scanner (Eckart & Ziegler, Berlin, Germany). Cyclone phosphor autoradiographic imaging system (Packard Instruments Co, New Jersey, USA) and Optiquant Imaging System software (version 5.0) was used for analysis. Immunostaining of brain sections were carried out by UCI Pathology core services. QuPath (version QuPath-0.2.2) was used for quantitative analysis of scanned brain slices.

### 2.2. Brain Tissue

Human postmortem brain tissue samples of AD, DSAD and CN cases (male and female), each including FCX and TCX were obtained from UCI Memory Impairment and Neurological Disorders (MIND) institute for in vitro experiments (Table 1). Brain slices, 10 µm thick, were obtained from frozen tissue on a Leica 1850 cryotome cooled to -20°C and collected on Fisher slides. All slides were then stored at -80°C. All postmortem human brain studies were approved by the Institutional Biosafety Committee of University of California, Irvine.

**Table 1.** Patient Case samples and data.

| ID<sup>1</sup> | Pathology | Gender | Age At Death | PMI<sup>2</sup> | Braak Score<sup>3</sup> | Plaque Stage<sup>4</sup> | Tangle Stage<sup>5</sup> |
| --- | --- | --- | --- | --- | --- | --- | --- |
| 01-25 | CN | Male | 83 | 1.8 | II | 0 | 2 |
| 02-11 | CN | Male | 96 | 3.58 | II | 0 | 2 |
| 03-07 | CN | Female | 84 | 4.25 | III | A | 3 |
| 12-09 | CN | Female | 95 | 2.92 | II | 0 | 2 |
| 17-14 | CN | Female | 89 | 5.95 | III | 0 | 3 |
| 06-18 | AD | Male | 46 | 3 | VI | C | 6 |
| 06-36 | AD | Male | 61 | 5.58 | VI | C | 6 |
| 12-26 | AD | Male | 55 | 3.18 | VI | C | 6 |
| 09-05 | AD | Female | 57 | 3.17 | VI | C | 6 |
| 16-38 | AD | Female | 76 | 3.82 | VI | B | 6 |
| 11-30 | DSAD | Male | 66 | 4.08 | VI | C | 6 |
| 08-42 | DSAD | Male | 55 | 4.5 | VI | C | 6 |
| 10-31 | DSAD | Female | 62 | 2.42 | VI | C | 6 |
| 07-31 | DSAD | Female | 52 | 4.37 | VI | C | 6 |
| 12-36 | DSAD | Female | 56 | 4.08 | VI | C | 6 |

Frozen brain samples of FCX and TCX were obtained from UCI MIND Institute; CN = cognitively normal; AD = Alzheimer's disease; DSAD= Down syndrome-associated Alzheimer's disease. <sup>1</sup>Each autopsy case identified by the year and order of autopsy (e.g. 08-42 is the 42nd autopsy performed in 2008). <sup>2</sup>PMI: Postmortem interval in hours. <sup>3</sup>Braak staging reflects pTau181 burden. <sup>4</sup>Plaque stage: Includes neuritic, cored and diffuse. Semi-quantitative scores of none, sparse, moderate and frequent were converted to a Plaque stage of A – C. <sup>5</sup>Tangle stage: neurofibrillary tangle density indicated by numerical values 0 – 6 for Tangle stage.

### 2.3. Immunohistochemistry

Adjacent brain slices were immunostained for tau and Aβ plaques. For total tau, DAKO polyclonal antibody detects all 6 six major isoforms of tau from the microtubule-associated protein tau gene and was used at a dilution 1:3000, A0024 (Agilent, CA, USA) using reported protocols (Ercan et al. 2017). Adjacent brain slices from all cases were immunostained with anti-Aβ Biolegend 803015 (Biolegend, CA, USA) which is reactive to amino acid residue 1-16 of β-amyloid. Anti-tau and anti-Aβ immunostained slides were scanned using the Ventana Roche slide scanner and the images generated were used for analysis by QuPath.

### 2.4. Autoradiography

#### 2.4.1. [^125^I]IPPI for Tau

Brain sections were treated with [^125^I]IPPI (60 mL; 3.7 kBq/mL, specific activity >90 GBq/µmol) in 10% ethanol phosphate buffer saline (PBS) buffer pH 7.4 (Syed et al. 2023). The chambers were incubated at 25°C for 1.25 hours and underwent multiple washes as reported [24]. The brain sections were air dried and transferred to a film cassette with a phosphor screen film inside that was exposed after 2 weeks. After removal from the cassette, the films were then read on the Phosphor Autoradiographic Imaging System/Cyclone Storage Phosphor System (Packard Instruments Co). Regions of interest (ROIs) were drawn on the autoradiographic images and the extent of binding of [^125^I]IPPI was measured in digital light units (DLU/mm^2^) using OptiQuant.

#### 2.4.2. [^125^I]IBETA for Aβ Plaques

Brain sections were treated with [^125^I]IBETA (60 mL; 5 kBq/mL, specific activity >90 GBq/µmol) using previously reported methods (Nguyen et al. 2022; Syed et al. 2023) . The brain sections were air dried and transferred to a film cassette with a phosphor screen film inside that was exposed overnight. The films were then removed from the film cassette and placed on the Phosphor Autoradiographic Imaging System/Cyclone Storage Phosphor System (Packard Instruments Co). ROIs were drawn on the autoradiographic images and the extent of binding of [^125^I]IBETA was measured using OptiQuant.

#### 2.4.3 [^125^I]IPPI In Vitro Drug Inhibition With Harmine

Inhibitor concentration (IC_50_) of harmine on [^125^I]IPPI binding was determined by using adjacent brain slices of FCX and TCX DSAD cases with different concentrations of harmine (10^-9^ to 10^-5^M). Unlabeled MK-6240 at 10 µM was designated for nonspecific binding to determine specific binding of [^125^I]IPPI. Binding studies used 3.7 kBq/mL of [^125^I]IPPI per cc of ethanol (20%) with incubation at 25°C for 75 minutes. Then the slices underwent two washes of 90% ethanol for 3 minutes each, one PBS buffer wash for 3 minutes and one cold water wash for 2 minutes. Slides of the brain sections were air dried and placed into a film cassette with a phosphor screen film. After two weeks the slides were removed from the cassette and the film was exposed and read by the Phosphor Autoradiographic Imaging System/Cyclone Storage Phosphor System (Packard Instruments Co). ROIs were drawn on the autoradiographic images and [^125^I]IPPI binding was measured in DLU/mm^2^ using the OptiQuant acquisition and analysis program (Packard Instruments Co). Nonspecific binding was subtracted from the total binding to calculate specific binding at different concentrations of harmine. Inhibitor concentration, IC_50_ was measured by plotting specific binding against harmine concentrations using GraphPad Prism 10.

#### 2.4.4 [^125^I]IPPI In Vitro Drug Inhibition With KuFal194

To investigate DYRK1A effects, KuFal194 (10-iodo-11H-indolo[3,2-c]quinoline-6-carboxylic acid) (Paclibar et al. 2025) was used on [^125^I]IPPI binding to adjacent slices of FCX and TCX DSAD cases. In our previous in vitro work in AD brain sections with [^125^I]KuFal194, binding was enhanced in the presence of urea, 1,3-bis(4-cyanophenyl)urea (BCU, 10 µM) [28]. Inclusion of BCU has been shown to assist in membrane permeability of carboxylic acid containing drugs like KuFal194 (Salam et al. 2021). Thus, the incubation of [^125^I]IPPI in the total binding and competition experiments with KuFal194 (10 µM) included the urea, BCU (10 µM).

### 2.5. Optiquant Image Analysis

ROIs were drawn on the autoradiographic images of FCX and TCX brain slices using Optiquant which gave measures of DLU/mm^2^ from the pixels of an autoradiographic image. Background activity levels were subtracted from all images. Higher DLU/mm^2^ from autoradiography indicated higher [^125^I]IBETA and [^125^I]IPPI binding.

### 2.6. QuPath Image Analysis

Using QuPath, annotations were made for Aβ plaques or tau in regions of the IHC brain slices of each case. Approximately 20-25 annotations were made for Aβ plaques or tau by means of visual identification. Negative annotations with no Aβ plaques or tau were drawn in each case. A pixel thresholder was created to outline the immunohistochemistry (IHC) images for Aβ plaques and tau separately. The pixel classifier was run on each entire brain slice and trained by the annotations made to generate a new image of the brain slice with pixels to indicate Aβ plaques or tau.

### 2.7. Statistical Analysis

GraphPad Prism 10 and Microsoft Excel 16 were used to assess DLU/mm^2^ values from OptiQuant. Statistical power was determined by performing student’s t-test in GraphPad Prism 10 and p values <0.05 indicated statistical significance. Error bars signify standard deviation within each group. The Shapiro-Wilk test confirmed normality of distribution among all groups except for [^125^I]IBETA FCX CN GM DLU/mm^2^ values. Parametric comparisons between [^125^I]IPPI and [^125^I]IBETA DLU/mm^2^ values within DSAD and AD cases were determined with Pearson’s correlation and linear regression in GraphPad Prism 10. The interquartile range method was used to determine any outliers within the datasets.

## 3. Results

### 3.1 [^125^I]IPPI Binding in DSAD, AD, and CN Cases

The binding of the radioiodinated tau imaging agent, [^125^I]IPPI, was evaluated in all DSAD cases (Figure 1). Figure 1A-D shows adjacent slices of representative case 08-42 DSAD FCX while Figure 1E-H shows case 08-42 DSAD TCX. The presence of various amounts of tau was confirmed by the anti-tau immunostainings of adjacent brain slices for all cases (Figure 1B & 1F). The tau pixel classifier confirmed the abundance of tau mostly within the GM regions (Figure 1C & 1G). There was an alignment between the amount of anti-tau detected and the [^125^I]IPPI binding on autoradiographic images. The [^125^I]IPPI binding for TCX (Figure 1I) and FCX (Figure 1J) cases displayed substantially more binding in GM compared to WM. There is significantly more binding in DSAD FCX GM than DSAD TCX GM among all DSAD cases (Figure 1K).

**Figure 1.**
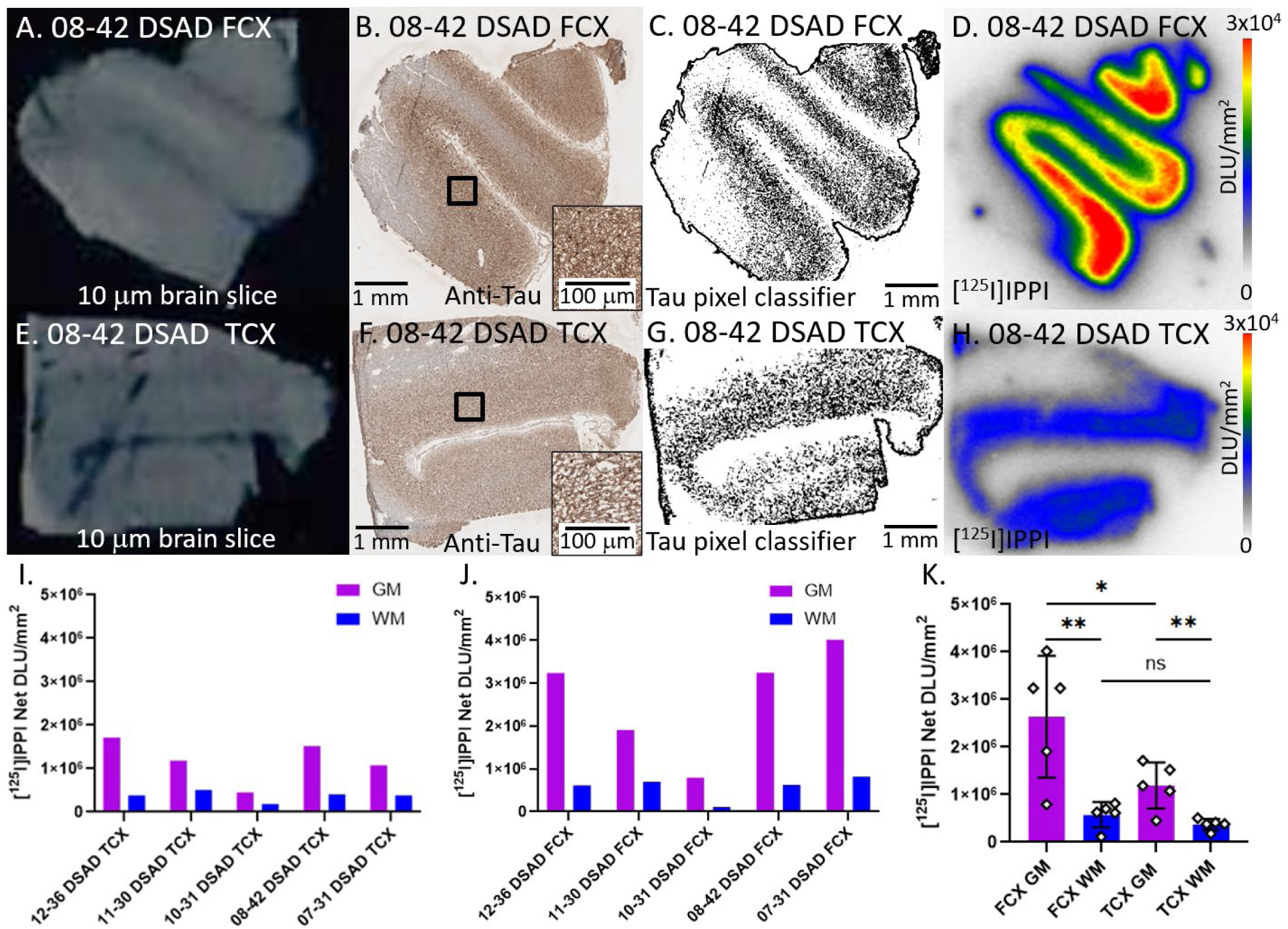
[^125^I]IPPI binding to Tau in DSAD: (A). Postmortem human brain slice (10 μm) of DSAD FCX 08-42; (B). Anti-tau IHC of adjacent brain slice of DSAD FCX 08-42 (1 mm magnification), inset of 100 µm magnification; (C). Tau pixel classifier image of DSAD FCX 08-42 (1 mm magnification); (D). [^125^I]IPPI binding to DSAD FCX 08-42 with little binding to WM regions; (E). Postmortem human brain slice (10 μm) of DSAD TCX 08-42; (F). Anti-tau IHC of adjacent brain slice of DSAD TCX 08-42 (1 mm magnification), inset of 100 µm magnification; (G). Tau pixel classifier image of DSAD TCX 08-42 (1 mm magnification); (H). [^125^I]IPPI binding to DSAD TCX 08-42 with little binding to WM regions; (I). [^125^I]IPPI binding to all DSAD TCX cases GM and WM regions; (J). [^125^I]IPPI binding to all DSAD FCX cases GM and WM regions; (K). Average [^125^I]IPPI binding to DSAD FCX and TCX cases GM and WM regions (* p < 0.05, ** p < 0.01, ns = not significant).

All AD cases were evaluated for [^125^I]IPPI binding (Figure 2). Figure 2A-D shows adjacent slices of representative case 12-26 AD FCX while Figure 2E-H shows case 12-26 AD TCX. Anti-tau immunostains (Figure 2B & 2F) of the five AD cases revealed the presence of tau with confirmation from the tau pixel classifier (Figure 2C & 2G). There was general alignment between the amount of anti-tau detected and the [^125^I]IPPI binding on AD TCX autoradiographic images. The [^125^I]IPPI binding for TCX (Figure 2I) and FCX (Figure 2J) cases showed greater binding in GM compared to WM. There was no significant difference in [^125^I]IPPI binding between AD FCX GM and AD TCX GM (Figure 2K).

**Figure 2.**
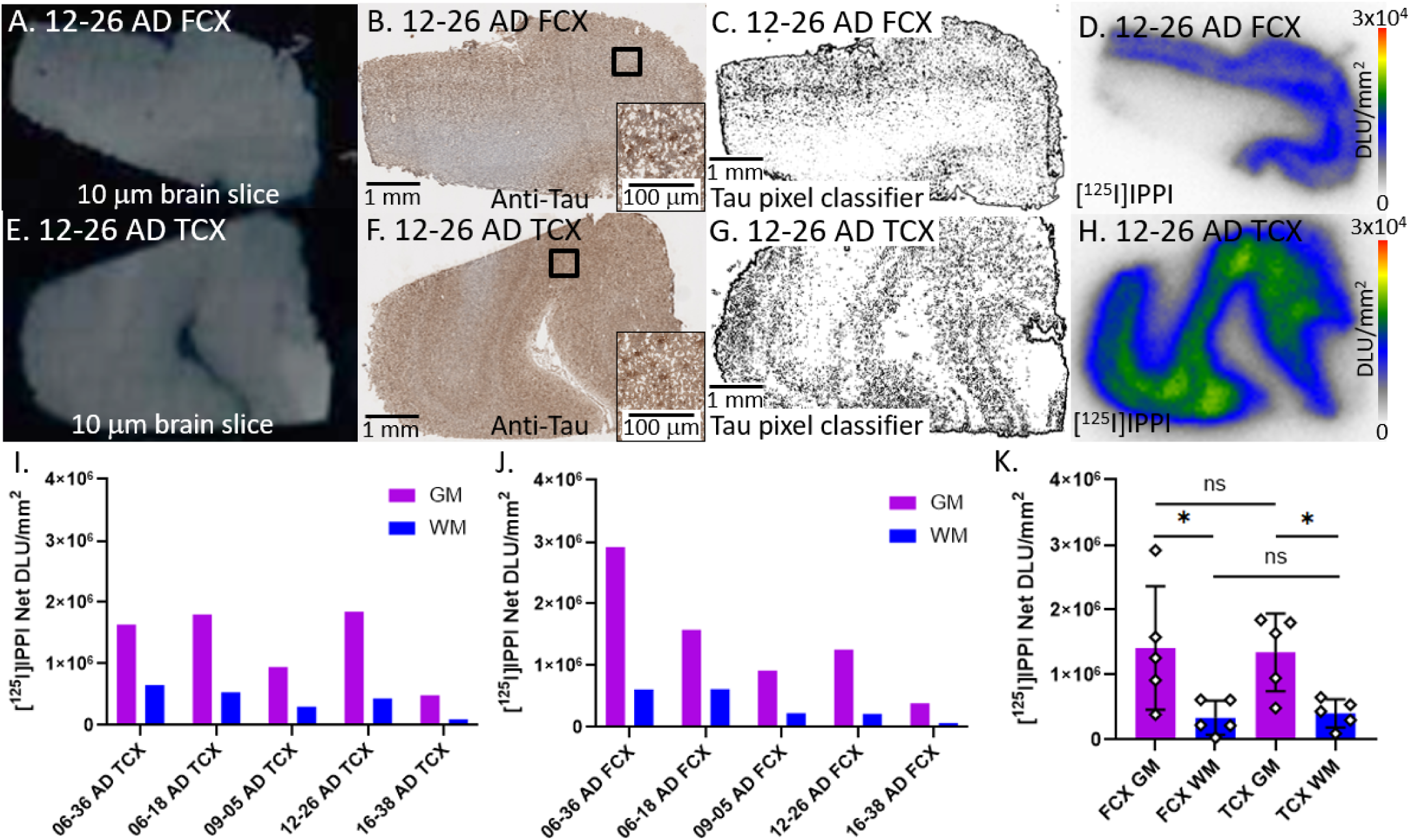
[^125^I]IPPI binding to Tau in AD: (A). Postmortem human brain slice (10 μm) of AD FCX 12-26; (B). Anti-tau IHC of adjacent brain slice of AD FCX 12-26 (1 mm magnification), inset of 100 µm magnification; (C). Tau pixel classifier image of AD FCX 12-26 (1 mm magnification); (D). [^125^I]IPPI binding to AD FCX 12-26 with little binding to WM regions; (E). Postmortem human brain slice (10 μm) of AD TCX 12-26; (F). Anti-tau IHC of adjacent brain slice of AD TCX 12-26 (1 mm magnification), inset of 100 µm magnification; (G). Tau pixel classifier image of AD TCX 12-26 (1 mm magnification); (H). [^125^I]IPPI binding to AD TCX 12-26 with little binding to WM regions; (I). [^125^I]IPPI binding to all AD TCX cases GM and WM regions; (J). [^125^I]IPPI binding to all AD FCX cases GM and WM regions; (K). Average [^125^I]IPPI binding to AD FCX and TCX cases GM and WM regions (* p<0.05, ns = not significant).

All CN cases were evaluated for [^125^I]IPPI binding (Figure 3). Figure 3A-D shows adjacent slices of representative case 03-07 CN FCX while Figure 3E-H shows case 03-07 CN TCX. Anti-tau immunostains (Figure 3B & 3F) of the five CN cases revealed minimal presence of tau with confirmation from the tau pixel classifier (Figure 3C & 3G). There was alignment between the amount of anti-tau detected and the [^125^I]IPPI binding on AD TCX autoradiographic image except for 03-07 CN FCX that displayed the highest [^125^I]IPPI binding (Figure 3D). Among all the CN cases, 03-07 and 17-14 were assigned the highest tangle stage which may be the reason for the higher [^125^I]IPPI binding (Table 1). Regardless, all [^125^I]IPPI binding values in CN cases were substantially lower than AD and DSAD. There were no significant differences between the binding in FCX and TCX (Figure 3K).

**Figure 3.**
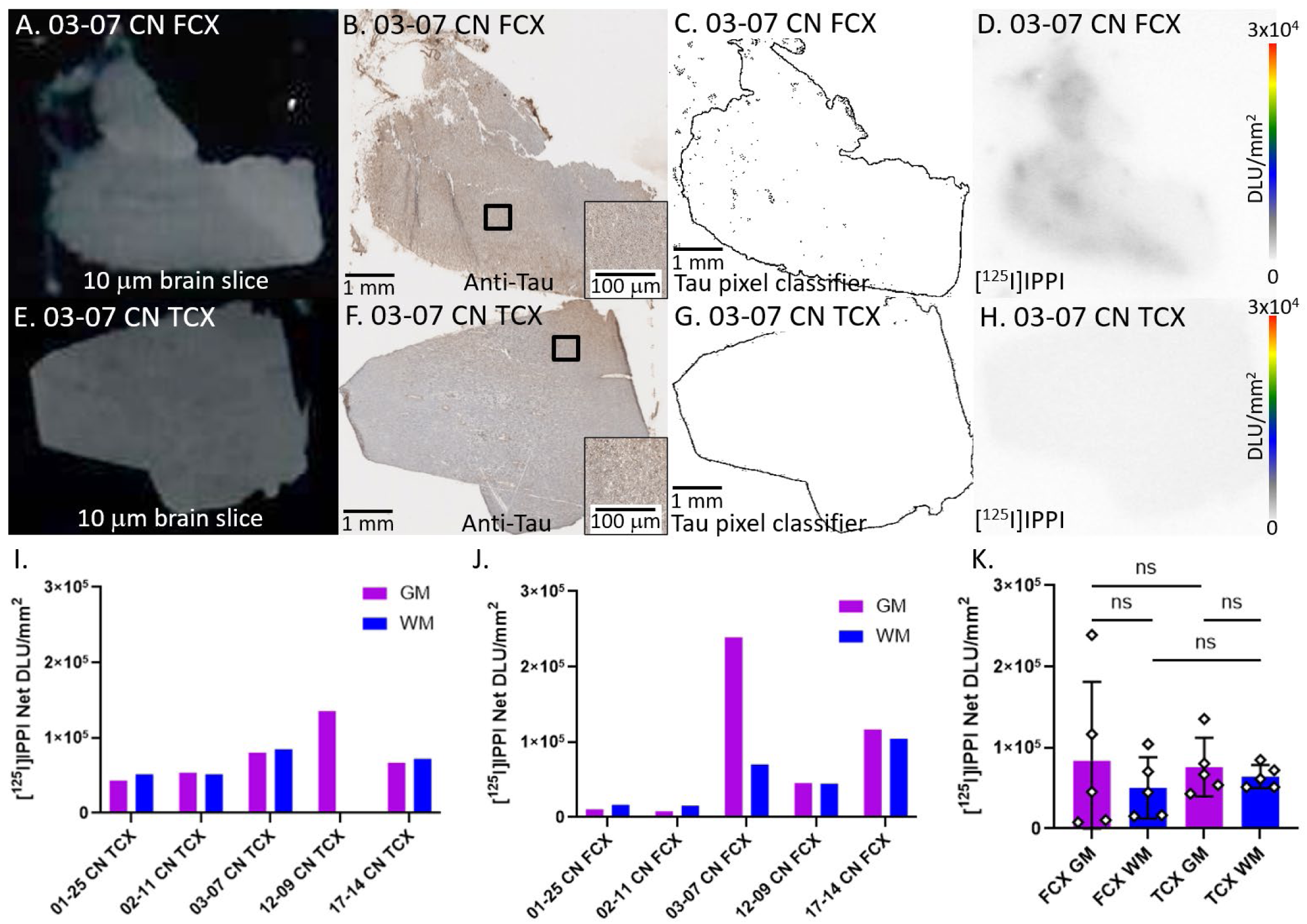
[^125^I]IPPI binding to Tau in CN: (A). Postmortem human brain slice (10 μm) of CN FCX 03-07; (B). Anti-tau IHC of adjacent brain slice of CN FCX 03-07 (1 mm magnification), inset of 100 µm magnification; (C). Tau pixel classifier image of CN FCX 03-07 (1 mm magnification); (D). [^125^I]IPPI binding to CN FCX 03-07 with little binding to WM regions; (E). Postmortem human brain slice (10 μm) of CN TCX 03-07; (F). Anti-tau IHC of adjacent brain slice of CN TCX 03-07 (1 mm magnification), inset of 100 µm magnification; (G). Tau pixel classifier image of CN TCX 03-07 (1 mm magnification); (H). [^125^I]IPPI binding to CN TCX 03-07 with little binding to WM regions; (I). [^125^I]IPPI binding to all CN TCX cases GM and WM regions; (J). [^125^I]IPPI binding to all AD FCX cases GM and WM regions; (K). Average [^125^I]IPPI binding to CN FCX and TCX cases GM and WM regions (ns = not significant).

### 3.2. In vitro Binding Inhibition of [^125^I]IPPI in DSAD

DSAD and AD demonstrate effective [^125^I]IPPI binding to FCX and TCX brain regions. To establish specificity of [^125^I]IPPI binding, drug competition studies were conducted using harmine that binds to MAO-A and DYRK1A [28]. Five concentrations of harmine were used on adjacent FCX and TCX brain slices. There were three adjacent brain slices for each case within each harmine concentration. MK-6240 was used on adjacent brain slices (Figure 4D) to obtain nonspecific binding to subtract from total binding of [^125^I]IPPI and [^125^I]IPPI + harmine binding at different concentrations to get specific binding. In Figure 4, representative case 08-42 DSAD FCX is shown in the absence and presence of harmine. Total binding (Figure 4A) was reduced in the presence of harmine (10^-7^ M visualized with lower [^125^I]IPPI binding, Figure 4B). As harmine concentrations increased, much of the bound [^125^I]IPPI was displaced (harmine 10^-5^ M, Figure 4C). This harmine competitive effect was consistent within both TCX and FCX regions and across different DSAD cases (Figure 4E). The measured average IC_50_ value for harmine was IC_50_ =115±40 nM across the three DSAD cases.

**Figure 4.**
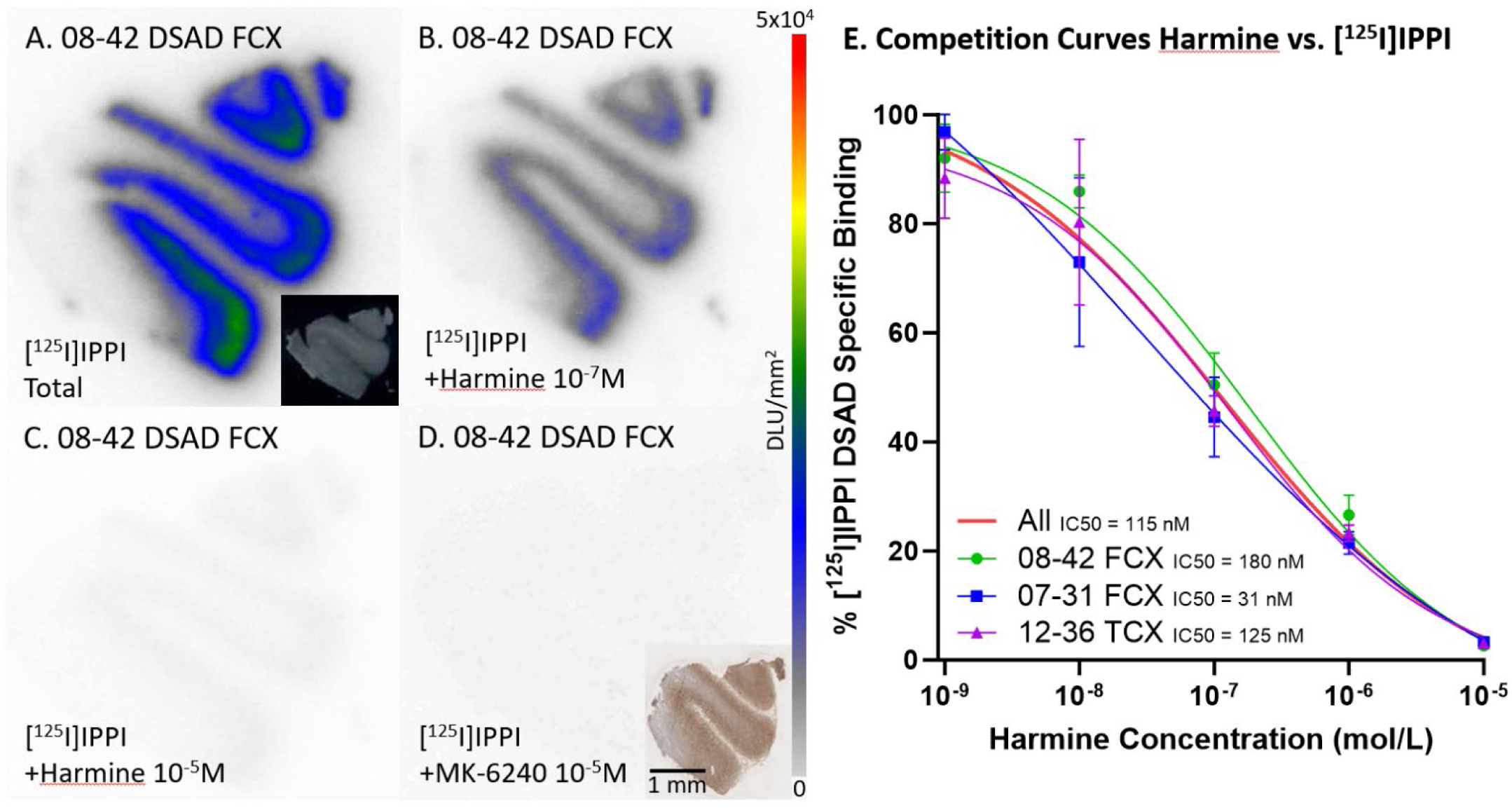
Harmine inhibition on [^125^I]IPPI Binding: (A). DSAD 08-42 FCX total [^125^I]IPPI binding without harmine; inset autoradiographic scan. (B). DSAD 08-42 FCX 10^-7^ harmine concentration on [^125^I]IPPI binding. (C). DSAD 08-42 FCX 10^-5^ harmine concentration on [^125^I]IPPI binding. (D). DSAD 08-42 FCX 10^-5^ MK-6240 concentration on [^125^I]IPPI binding; inset tau immunostaining of adjacent section (1 mm magnification). (E). Average [^125^I]IPPI specific binding of each brain region throughout all harmine concentrations. The red line represents the best fit line for all cases. Each brain region is plotted with one representative case with 3 brain slices for each concentration.

Since harmine binds to DYRK1A and inhibits [^125^I]IPPI binding, there was interest in observing if KuFal194 also inhibits [^125^I]IPPI binding. Structural similarities between IPPI (Figure 5A) and MK-6240 (Figure 5B) allow for both to bind to tau and thus MK-6240 inhibits [^125^I]IPPI binding. Although there are structural differences between harmine (Figure 5C) and KuFal194 (Figure 5D), they both can target DYRK1A. Among all the DSAD cases and both FCX and TCX regions, there was a small non-significant decrease in [^125^I]IPPI binding compared to total [^125^I]IPPI binding (Figure 5E).

**Figure 5.**
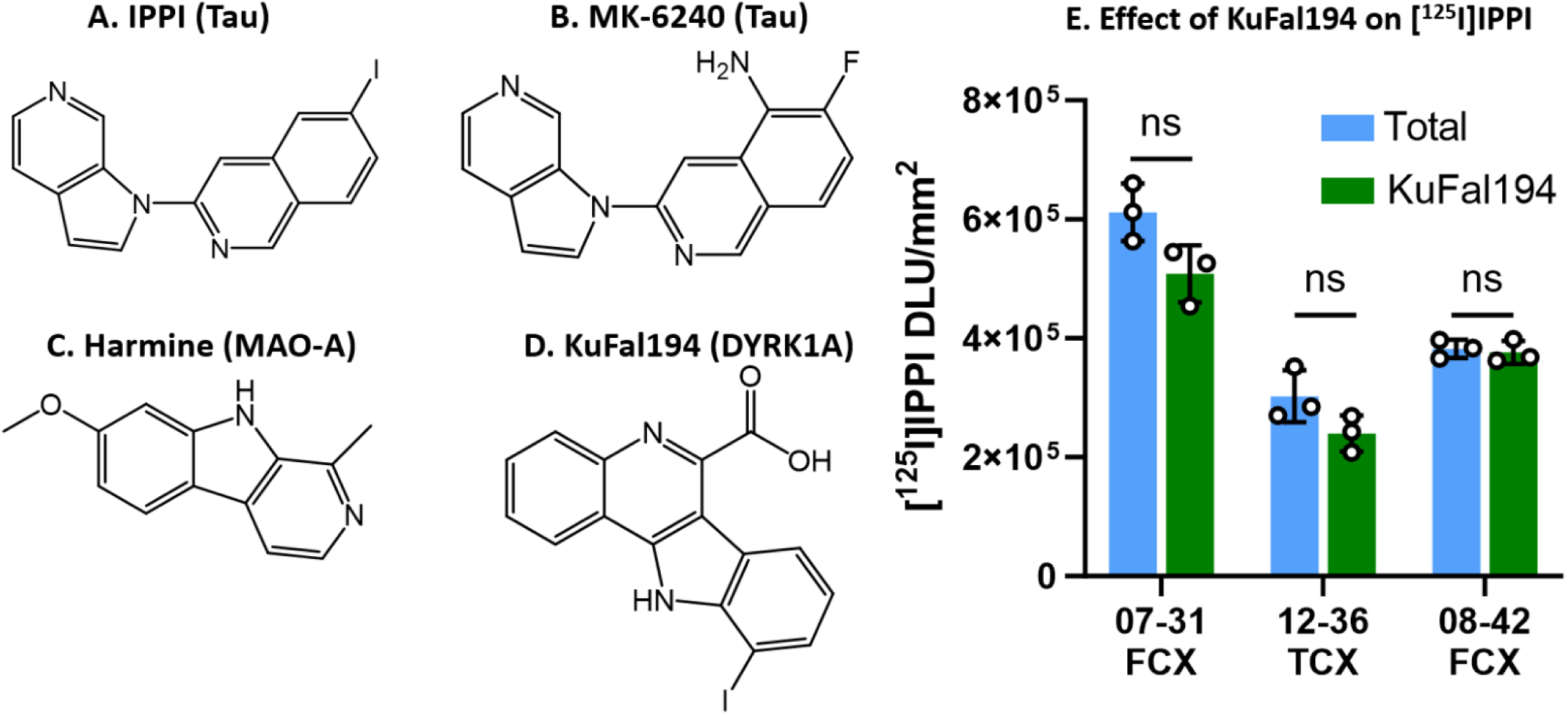
Effect of DYRK1A on [^125^I]IPPI Binding: Chemical structures of (A). IPPI; (B). MK-6240; (C). Harmine; (D). KuFal194; (E). Comparisons between total [^125^I]IPPI binding and [^125^I]IPPI binding with 10 μM KuFal194 on DSAD cases. All p-values for the t-tests were not significant (ns).

### 3.3. [^125^I]IBETA Binding in DSAD, AD, and CN Cases

The binding of the radioiodinated Aβ imaging agent, [^125^I]IBETA, was evaluated in all DSAD cases (Figure 6). Anti-Aβ immunostains of the five DSAD cases revealed the presence of extensive Aβ plaques (Figure 6C & 6D) especially in GM regions, confirmed by their respective Aβ pixel classifiers. This distribution aligned with the [^125^I]IBETA binding on autoradiographic images (Figure 6D & 6H). All cases displayed higher [^125^I]IBETA binding in GM compared to WM within both TCX (Figure 6I) and FCX (Figure 6J) regions. There were no significant differences in [^125^I]IBETA binding between FCX and TCX.

**Figure 6.**
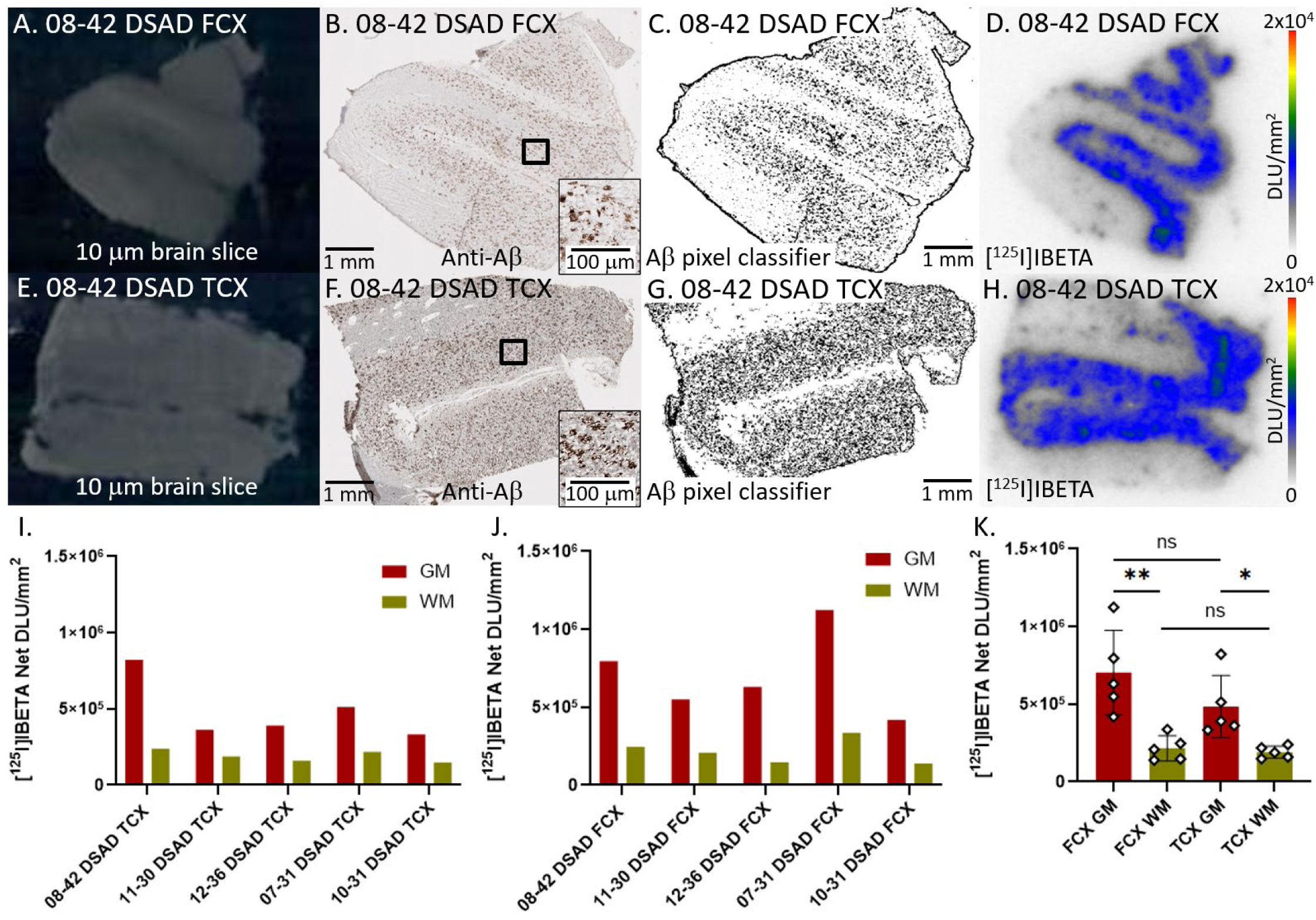
[^125^I]IBETA binding to Aβ plaques in DSAD: (A). Postmortem human brain slice (10 μm) of DSAD FCX 08-42; (B). Anti-Aβ IHC of adjacent brain slice of DSAD FCX 08-42 (1 mm magnification), inset of 100 µm magnification; (C). Aβ pixel classifier image of DSAD FCX 08-42 (1 mm magnification); (D). [^125^I]IBETA binding to DSAD FCX 08-42 with little binding to WM regions; (E). Postmortem human brain slice (10 μm) of DSAD TCX 08-42; (F). Anti-Aβ IHC of adjacent brain slice of DSAD TCX 08-42 (1 mm magnification), inset of 100 µm magnification; (G). Aβ pixel classifier image of DSAD TCX 08-42 (1 mm magnification); (H). [^125^I]IBETA binding to DSAD TCX 08-42 with little binding to WM regions; (I). [^125^I]IBETA binding to all DSAD TCX cases GM and WM regions; (J). [^125^I]IBETA binding to all DSAD FCX cases GM and WM regions; (K). Average [^125^I]IBETA binding to DSAD FCX and TCX cases GM and WM regions (* < 0.05, ** p < 0.01, ns = not significant).

All AD cases were evaluated for [^125^I]IBETA binding (Figure 7). Figure 7A-D showed adjacent slices of representative case 12-26 AD FCX while Figure 7E-H showed case 12-26 AD TCX. The presence of Aβ plaques was confirmed in the anti-Aβ immunostains of all the AD cases with the Aβ pixel classifiers visualizing Aβ distribution (Figure 7C & 7H). The Aβ distribution mostly aligned with the [^125^I]IBETA binding distribution in the postmortem brain slices (Figure 7D & 7H). There was higher [^125^I]IBETA binding in GM than WM regions among both TCX and FCX slices (Figure 7I & 7J). There was significantly more [^125^I]IBETA binding in FCX than TCX GM (Figure 7K).

**Figure 7.**
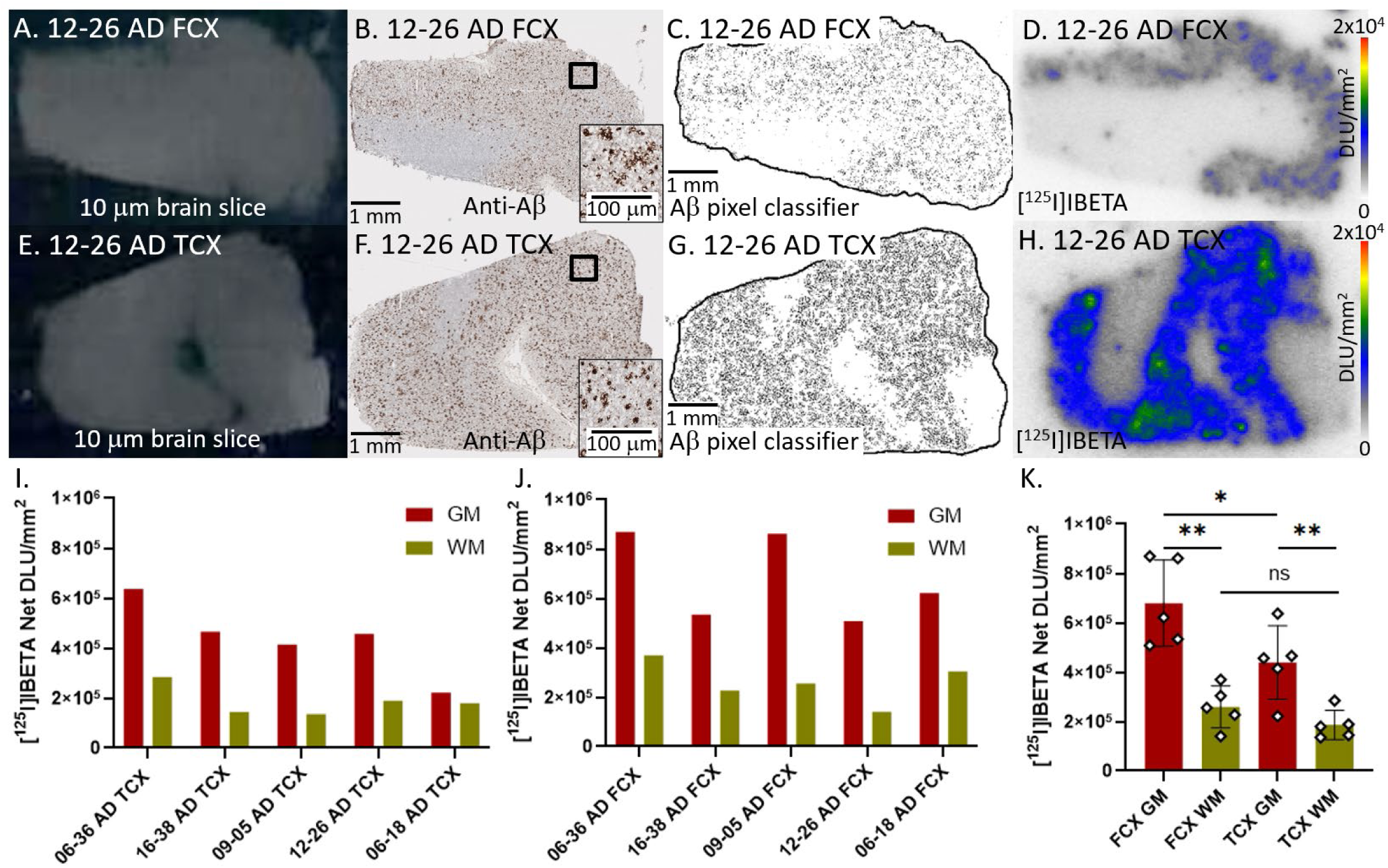
[^125^I]IBETA binding to Aβ plaques in AD: (A). Postmortem human brain slice (10 μm) of AD FCX 12-26; (B). Anti-Aβ IHC of adjacent brain slice of AD FCX 12-26 (1 mm magnification), inset of 100 µm magnification; (C). Aβ pixel classifier image of AD FCX 12-26 (1 mm magnification); (D). [^125^I]IBETA binding to AD FCX 12-26 with little binding to WM regions; (E). Postmortem human brain slice (10 μm) of AD TCX 12-26; (F). Anti-Aβ IHC of adjacent brain slice of AD TCX 12-26 (1 mm magnification), inset of 100 µm magnification; (G). Aβ pixel classifier image of AD TCX 12-26 (1 mm magnification); (H). [^125^I]IBETA binding to AD TCX 12-26 with little binding to WM regions; (I). [^125^I]IBETA binding to all AD TCX cases GM and WM regions; (J). [^125^I]IBETA binding to all AD FCX cases GM and WM regions; (K). Average [^125^I]IBETA binding to AD FCX and TCX cases GM and WM regions (* p < 0.05, ** p < 0.01, ns = not significant).

All five CN cases were assessed for [^125^I]IBETA binding. Figure 8A-D shows representative case 03-07 FCX while Figure 8E-H showed case 03-07 TCX. Anti-Aβ immunostains (Figure 8B & 8F) of all the CN cases demonstrate the lack of Aβ as confirmed by the Aβ pixel classifier (Figure 8C & 8G). However, 03-07 CN FCX exhibits some sparse Aβ (Figure 8C) that [^125^I]IBETA bound to (Figure 8D). This is the only CN case with a plaque score greater than 0 but this amount remains lower than all the AD cases (Table 1). The [^125^I]IBETA FCX CN GM DLU/mm^2^ values did not pass the Shapiro-Wilk test, confirming there is no normality of distribution within this dataset. The interquartile range method was performed on the dataset, indicating the value for 03-07 CN FCX as an outlier. When excluding the outlier 03-07 FCX, there were no significant differences between the [^125^I]IBETA binding in FCX and TCX regions (Figure 8K).

**Figure 8.**
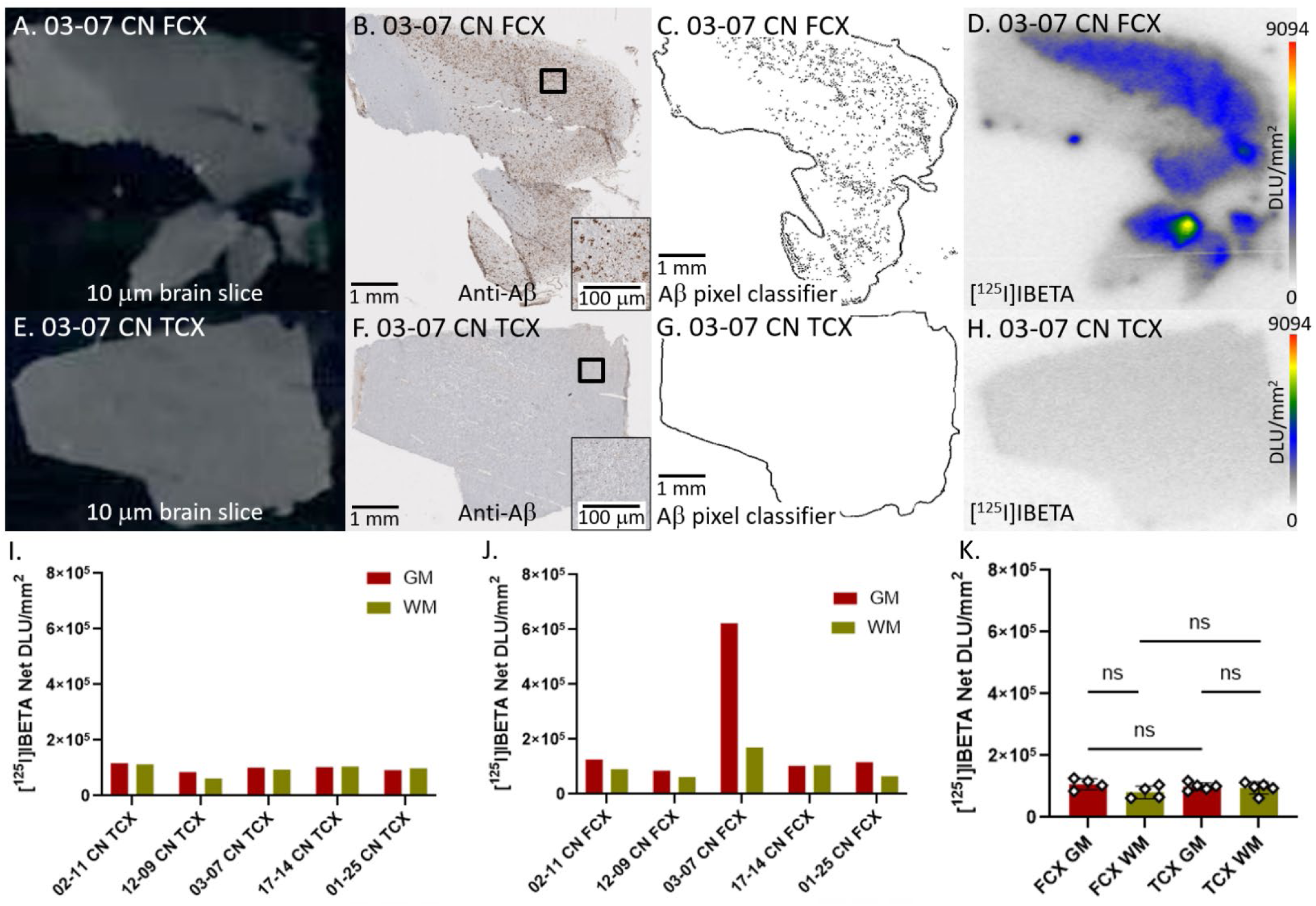
[^125^I]IBETA binding to Aβ plaques in CN: (A). Postmortem human brain slice (10 μm) of CN FCX 03-07; (B). Anti-Aβ IHC of adjacent brain slice of CN FCX 03-07 (1 mm magnification), inset of 100 µm magnification; (C). Aβ pixel classifier image of CN FCX 03-07 (1 mm magnification); (D). [^125^I]IBETA binding to CN FCX 03-07 with little binding to WM regions; (E). Postmortem human brain slice (10 μm) of CN TCX 03-07; (F). Anti-Aβ IHC of adjacent brain slice of CN TCX 03-07 (1 mm magnification), inset of 100 µm magnification; (G). Aβ pixel classifier image of CN TCX 03-07 (1 mm magnification); (H). [^125^I]IBETA binding to CN TCX 03-07 with little binding to WM regions; (I). [^125^I]IBETA binding to all CN TCX cases GM and WM regions; (J). [^125^I]IBETA binding to all AD FCX cases GM and WM regions; (K). Average [^125^I]IBETA binding to CN FCX and TCX cases GM and WM regions (ns = not significant).

### 3.4. [^125^I]IPPI and [^125^I]IBETA Binding Comparisons

Figure 9 presents overall comparisons between DSAD, AD, and CN GM and WM in TCX and FCX [^125^I]IPPI and [^125^I]IBETA binding. The amount of radiotracer binding to GM and WM in DSAD and AD cases remained significantly higher than CN throughout FCX (Figure 9B, 9D) and TCX (Figure 9A, 9C) regions. However, there were no significant differences between DSAD and AD in radiotracer binding to GM or WM. Although not significant, the greatest difference between DSAD and AD cases was 48% within [^125^I]IPPI binding to FCX GM (Figure 9B).

**Figure 9.**
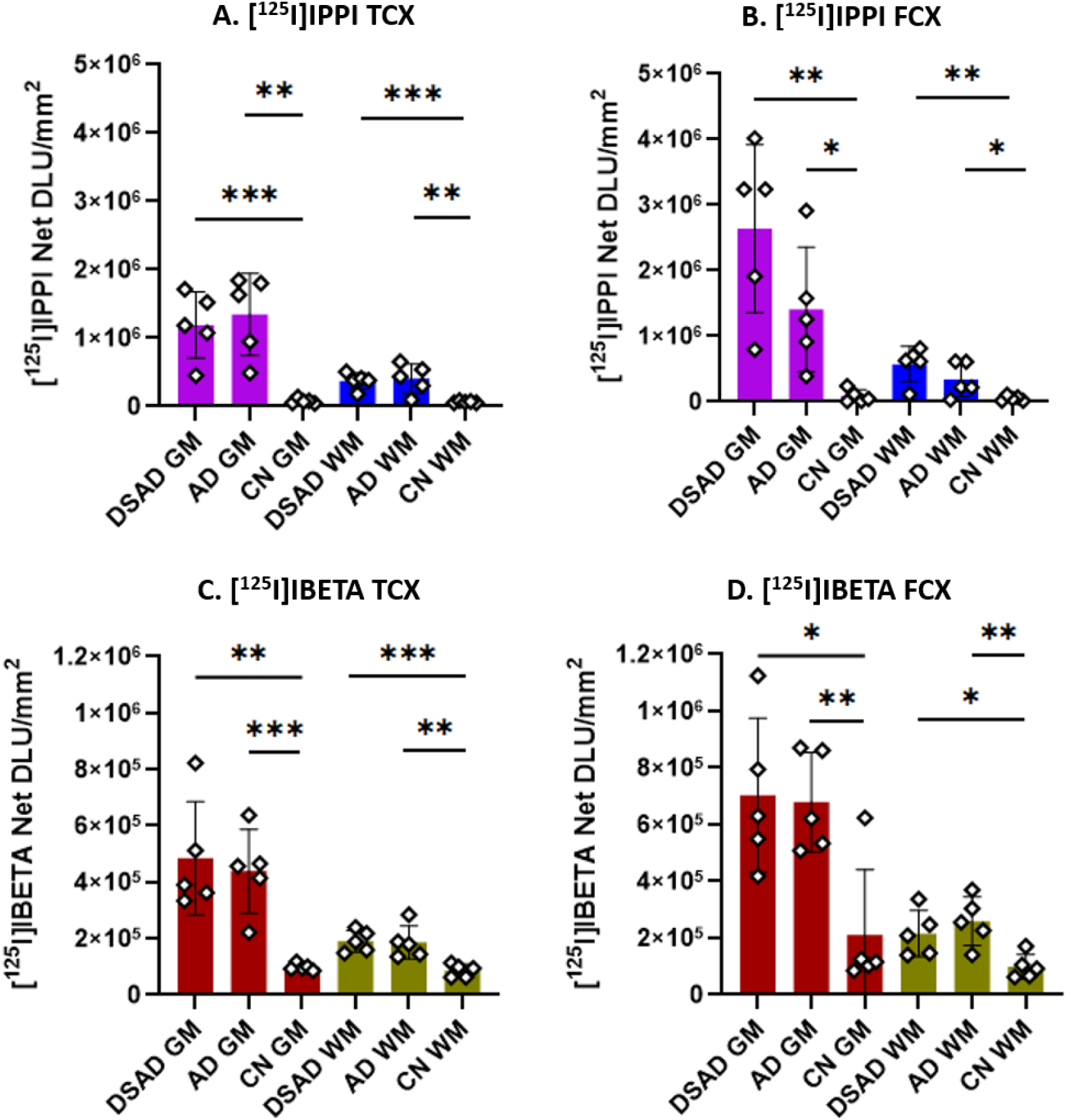
Group Comparisons of [^125^I]IPPI and [^125^I]IBETA binding in GM and WM: Unpaired two-tailed parametric t-tests determined statistical significance between each parameter (* p < 0.05, ** p < 0.01, *** < 0.001, ns = not significant). (A). [^125^I]IPPI binding in TCX GM and WM of DSAD, AD, and CN cases. (B). [^125^I]IPPI binding in FCX GM and WM of DSAD, AD, and CN cases. (C). [^125^I]IBETA binding in TCX GM and WM of DSAD, AD, and CN cases. (D). [^125^I]IBETA binding in FCX GM and WM of DSAD, AD, and CN cases.

To observe the contrast between GM and WM binding of the radiotracers, GM/WM ratios were determined within each case for FCX and TCX. The GM/WM ratios for both [^125^I]IPPI (Figure 10A) and [^125^I]IBETA (Figure 10B) varied in significance. The average [^125^I]IPPI GM/WM ratios of DSAD FCX (5.0), DSAD TCX (3.2), AD FCX (4.7), and AD TCX (3.7) were significantly greater than CN FCX (1.3) and CN TCX (1.2) (Figure 10A). Similarly, average [^125^I]IBETA GM/WM ratios of DSAD FCX (3.3), DSAD TCX (2.5), AD FCX (2.7), and AD TCX (2.4) were significantly greater than CN FCX (1.4) and CN TCX (1.1) (Figure 10B). The average GM/WM ratios of all DSAD cases ([^125^I]IPPI=4.1, [^125^I]IBETA=2.9) and all AD cases ([^125^I]IPPI=4.2, [^125^I]IBETA=2.6) were significantly greater than CN cases ([^125^I]IPPI=1.3, [^125^I]IBETA=1.2). Overall, FCX GM/WM ratios were greater than TCX GM/WM ratios among all groups.

**Figure 10.**
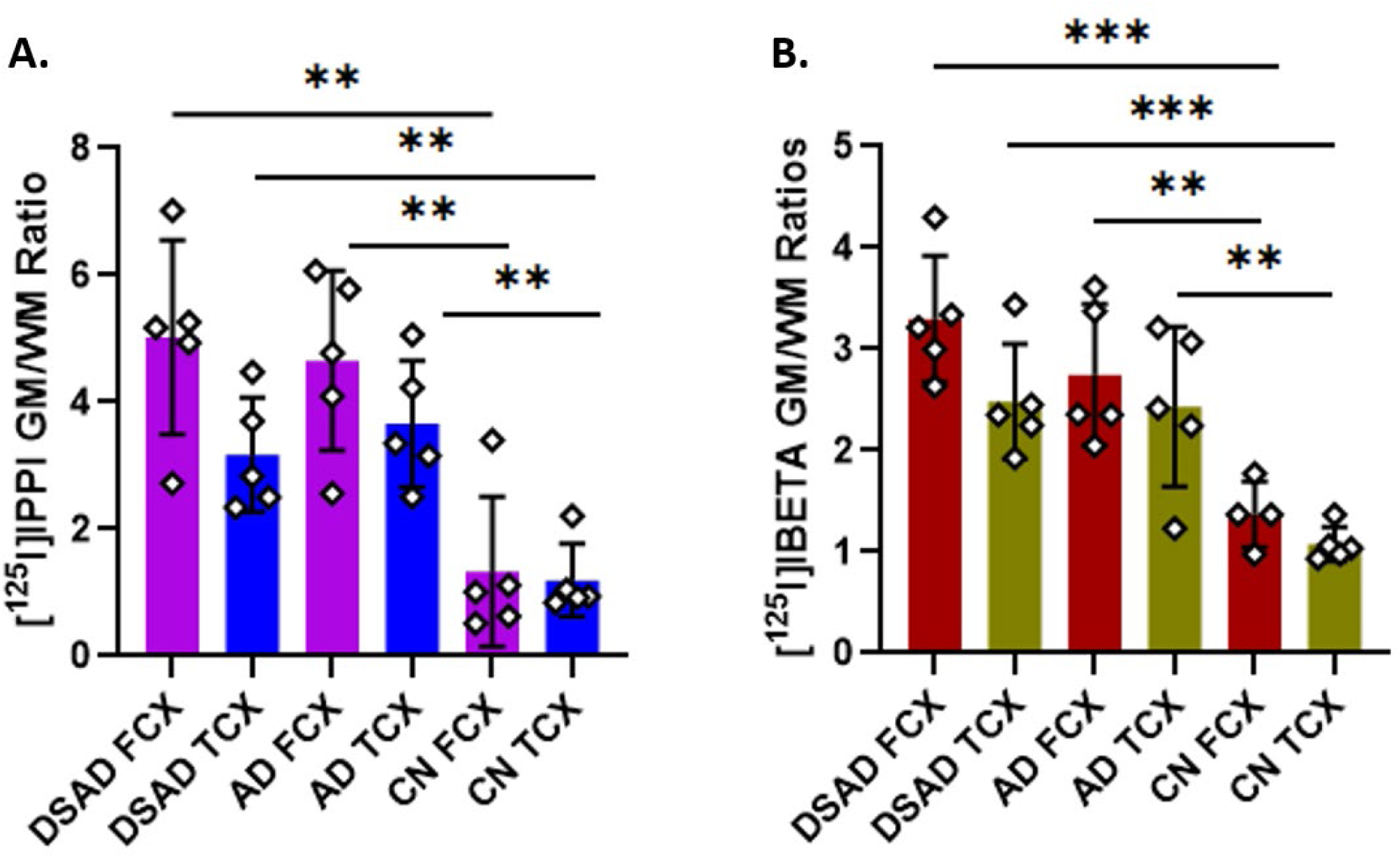
Group Comparisons of [^125^I]IPPI and [^125^I]IBETA GM/WM Ratios: Unpaired two-tailed parametric t-tests determined statistical significance between each parameter (* p < 0.05, ** p < 0.01, *** < 0.001, ns = not significant). (A). [^125^I]IPPI GM/WM ratios of FCX and TCX in DSAD, AD, and CN cases. (B). [^125^I]IBETA GM/WM ratios of FCX and TCX in DSAD, AD, and CN cases.

The relationship between [^125^I]IPPI and [^125^I]IBETA binding was evaluated by correlating the two radiotracers (Figure 11). The datasets of [^125^I]IPPI and [^125^I]IBETA binding were individually tested and confirmed for normality of distribution using the Shapiro-Wilk test. With this, Pearson’s correlation was done to determine the parametric correlation between [^125^I]IPPI and [^125^I]IBETA binding. Both DSAD (Figure 11A) and AD (Figure 11B) cases showed a positive correlation between [^125^I]IPPI and [^125^I]IBETA binding. DSAD cases demonstrated a stronger correlation (Pearson’s *r* = 0.4528) than AD cases (Pearson’s *r* = 0.4274).

**Figure 11.**
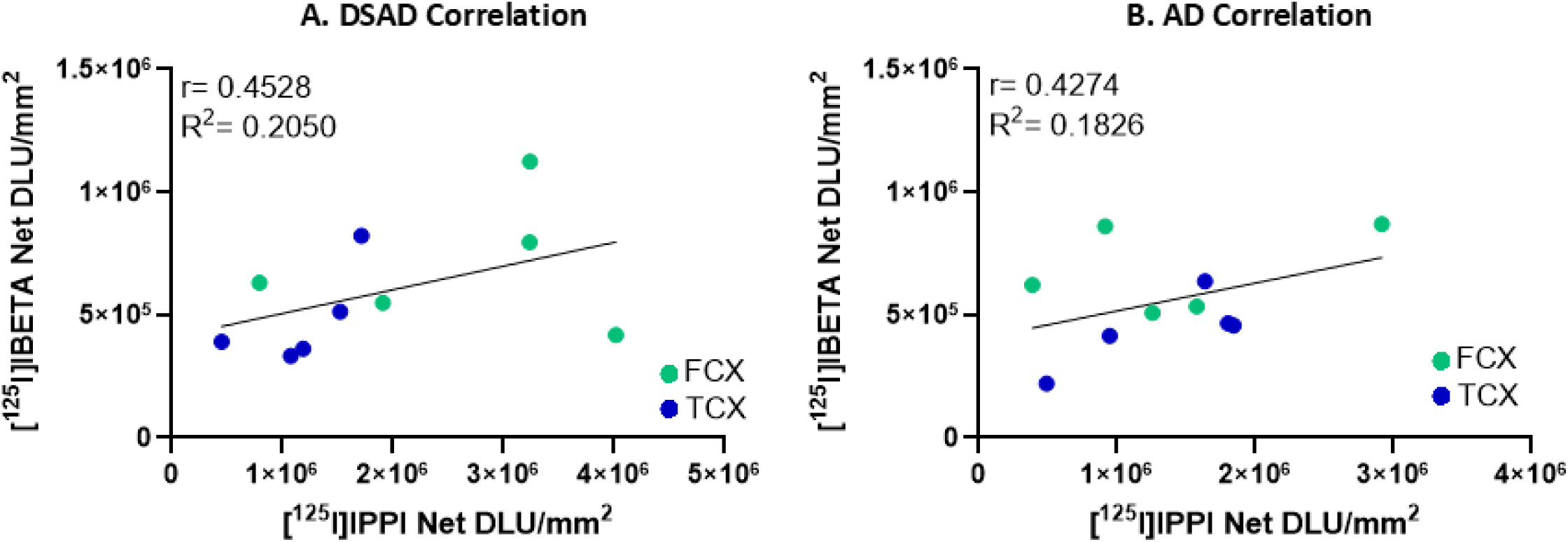
Correlation of [^125^I]IPPI and [^125^I]IBETA binding to tau and Aβ plaque respectively: (A). [^125^I]IPPI and [^125^I]IBETA binding in DSAD cases (Pearson’s *r*=0.4528; R^2^=0.2050); (B). [^125^I]IPPI and [^125^I]IBETA binding in AD cases (Pearson’s *r*=0.4274; R^2^=0.1826).

## 4. Discussion

As molecular biomarkers, tau and Aβ have become indispensable for assisting in the clinical diagnosis of AD and are now being applied to study AD pathogenesis in people with DS, emphasizing a continued need to develop PET imaging agents (Chrem Mendez et al. 2019). In this study, the binding of [^125^I]IPPI and [^125^I]IBETA to tau and Aβ plaques respectively in FCX and TCX regions was highly selective in DSAD and AD cases. The extent of overall binding of the two radiotracers was similar or greater in DSAD cases compared to the AD cases. The CN cases showed minimal binding as expected. The GM/WM ratios averaged higher in FCX than TCX. There was a positive correlation between [^125^I]IPPI and [^125^I]IBETA binding.

As expected, all DSAD and AD cases exhibited greater [^125^I]IPPI and [^125^I]IBETA binding than CN cases where all GM comparisons were significantly different in both FCX and TCX while most WM comparisons were significant (Figure 9). Although there was one CN case (03-07) with higher [^125^I]IPPI and [^125^I]IBETA binding than the rest of the CN cases in the FCX, this is confirmed by the higher tau and Aβ scores (Table 1). The assigned scores and binding remained less than DSAD and AD cases, but this case may indicate prodromal AD. All DSAD and AD cases were Braak stage VI while CN 03-07 was Braak stage III, in addition to being the only case with a plaque stage of A, contributing to the exacerbated difference. Relative to CN cases, previous studies suggest that DS FCX show high amounts of soluble Aβ40 and Aβ42 and higher tau phosphorylation (Di Domenico et al. 2013). In plaques, Aβ42 precedes Aβ40 plaque formation in DS, so its abundance in CN cases indicate the severity of neurodegeneration that can soon lead to AD or DSAD (Teller et al. 1996; Di Domenico et al. 2013).

In FCX and TCX brain slices, all DSAD and AD cases observed greater [^125^I]IPPI and [^125^I]IBETA binding in GM than WM. Chromosome 21 is triplicated in DS and contains an amyloid precursor protein that is cleaved in the amyloidogenic pathway, resulting in greater brain Aβ levels with increasing DS age (Hampel et al. 2021). The progression of Aβ plaque accumulation occurs within the same brain regions as AD, soon reaching sufficient levels for an AD diagnosis. Interestingly, the striatum was the first region to show elevated [^11^C]PiB retention in DS followed by anterior cingulate, FCX, and TCX despite the striatum typically being spared until later AD pathogenesis (Lao et al. 2016). The distribution of [^11^C]PiB binding in particular brain regions may reveal differences in Aβ plaque severity between DSAD and AD. In DSAD cases, [^125^I]IPPI binding to tau was significantly greater in FCX than TCX but this significance was not observed in DSAD [^125^I]IBETA binding. The onset and progression of tau and Aβ plaque pathology have been regarded as similar between GM TCX and FCX in DSAD brains (Abrahamson et al. 2019). Tau and Aβ plaque pathology have also been suggested to be more prevalent in DSAD FCX due to significantly higher [^11^C]-PIB and ^3^H-THK5117 binding compared to CN (Lemoine et al. 2020). Most literature focuses on the prevalence of FCX rather than TCX so more comprehensive comparisons between various brain regions are needed to ascertain the progression of tau and Aβ plaque pathology in DSAD.

To expand on the specificity of [^125^I]IPPI binding to tau in FCX and TCX DSAD brain regions, it is essential to test how drugs that bind to tau related proteins affect [^125^I]IPPI binding. Given that MK-6240 and IPPI share structural similarities, MK-6240 greatly inhibits [^125^I]IPPI binding and was therefore used for nonspecific binding (Figure 4D). Harmine significantly inhibited [^125^I]IPPI binding with increasing concentrations and the IC_50_ were similar between TCX and FCX, indicating similar inhibition patterns (Figure 4E). Harmine (a known MAO-A and DYRK1A inhibitor) (Paclibar et al. 2025) has previously been shown to substantially displace the tau PET tracers [^18^F]flortaucipir and [^18^F]MK-6240 in AD brains (Aguero et al. 2024). Because of the inconsistent effect of clorgyline (MAO-A inhibitor) and harmine on [^18^F]MK-6240, it was suggested perhaps the effect of harmine on [^18^F]MK-6240 may be due to its binding to DYRK1A (Aguero et al. 2024). However, we have recently reported that harmine has a moderate affinity to tau and therefore displaces tau-binding drugs, including [^125^I]IPPI binding in AD brain slices (Karim et al. 2025). Therefore, harmine has multiple targets which include MAO-A, DYRK1A and tau with a binding affinity order of MAO-A (5nM) > DYRK1A (70 nM) > tau (135 nM) (Paclibar et al. 2025; Karim et al. 2025). DYRK1A resides on chromosome 21, thus DS involves the overexpression of DYRK1A that contributes to several phenotypes associated with DS and alters neurological functions (Liu et al. 2008). DYRK1A can phosphorylate tau and potentially drive NFT associated with AD pathogenesis. Harmine is a potent DYRK1A inhibitor that has exemplified its ability to interfere with neurite formation and DYRK1A-induced tau phosphorylation (Göckler et al. 2009; Frost et al. 2011). Compared to the effect of harmine on [^125^I]IPPI in the same AD brains [35], harmine had a similar effect on [^125^I]IPPI in DSAD brains (IC_50_ =115±40 nM). To elucidate how harmine may have inhibited [^125^I]IPPI binding in the DSAD brains, KuFal194 binding to DYRK1A was observed but KuFal194 did not significantly inhibit [^125^I]IPPI binding in either FCX or TCX (Figure 5E). The minimal displacement of [^125^I]IPPI binding by KuFal194 may have been to tau within DYRK1A but other tau binding remained unaffected. Independently of DYRK1A, harmine binding significantly inhibits multiple forms of phosphorylated tau including total tau, both of which are increased in AD (Frost et al. 2011; Sjögren et al. 2001). These results further emphasize the unique nature of harmine in inhibiting the binding of tau radiotracers (albeit weakly) in addition to binding to DYRK1A associated with the formation of tau tangles.

As prominent AD neuropathologies, investigating potential relationships between tau and Aβ accumulation can provide valuable insight into disease progression. Within DSAD and AD cases, the positive correlation between [^125^I]IPPI binding to tau and [^125^I]IBETA binding to Aβ plaque indicates that their accumulation may be linked to one another. A synergistic relationship between tau and Aβ plaque has previously been suggested where both contribute to neurodegeneration, although the mechanism is not clearly understood (Grigorova et al. 2022). Tau and Aβ become toxic through independent but converging mechanisms but may amplify each other’s toxic effects where increasing tau concentrations in dendrites can make neurons more vulnerable to damage induced by Aβ (Ittner et al. 2011). These potential connections have been investigated within the context of AD but there may be a similar synergistic connection between tau and Aβ in DSAD pathology where they potentiate cognitive decline together. This idea was supported by the commonly observed pattern of amyloid deposition arising early in DS brains before tau pathology, but disease progression accelerates once both are present (Davidson et al. 2018; Condello et al. 2022). Thus, there appears to be a relationship between the two biomarkers, but this relationship needs to be investigated more thoroughly.

The small number of DSAD and AD cases with the same Braak stages is a limitation of this study along with some inter-case variability. Although the amount of [^125^I]IPPI and [^125^I]IBETA binding were consistent throughout all adjacent slices per case, minimizing inter-case variability would further validate the generality of these findings. The CN cases were older than both DSAD and AD cases, possibly indicating more radioligand binding than anticipated due to factors such as old age. The exclusion of outliers from a small sample size may skew certain comparisons which would otherwise not be a concern in a larger sample size. Therefore, these findings suggest potential in the radioligands [^125^I]IPPI and [^125^I]IBETA to be confirmed on a larger scale. To add onto these preliminary findings, future studies will include more cases with earlier progression of DSAD and AD with consideration for mild cognitive impairment. This will allow for translatable results comparative to the general patient population and more accurate correlation of Aβ plaque and tau positivity with [^125^I]IBETA and [^125^I]IPPI. Despite these limitations, the findings of this study establish [^125^I]IBETA and [^125^I]IPPI as promising radioligands in DSAD and AD. Future studies will aim to further investigate the effectiveness of [^125^I]IBETA and [^125^I]IPPI in other brain regions with more diverse cases and seek how to incorporate these radioligands as SPECT imaging analogs for in vivo diagnostic studies for DSAD and AD (Mondal et al. 2023).

## 5. Conclusions

Overall, the autoradiography binding profile of [^125^I]IBETA and [^125^I]IPPI to Aβ plaques and tau demonstrates their effectiveness as radioligands in both DSAD and AD brains. Binding profiles of these two radiotracers in DSAD and AD suggests similarity in effectiveness of these two biomarkers and thus allows a more comprehensive analysis of Aβ plaques and tau to reveal signs of disease progression and severity. Expanding upon techniques to evaluate Aβ plaques and tau in vitro enhances current understandings of the unique features of DSAD that may differ from AD and how to approach them. Thoroughly investigating the capabilities of tau and Aβ plaque radioligands is essential to elucidate how these biomarkers accumulate and contribute to DSAD and AD progression, eventually in vivo as well.

## Author Contributions

All authors had full access to all the data in the study and take responsibility for the integrity of the data and the accuracy of the data analysis. Study concept and design: JM. Acquisition of data: APB, FK, DMS, SAS, EH, CL and JM. Analysis and interpretation of data: APB, FK, DMS, SAS and JM. Drafting of the manuscript: FK, APB, and JM Statistical analysis: APB, FK. Obtained funding: JM. Study supervision: JM.

## Institutional Review Board Statement

Not applicable

## Informed Consent Statement

Not applicable.

## Data Availability Statement

The data that support the findings of this study are available from the corresponding author upon reasonable request.

## Acknowledgments

Research support provided by NIH AG R01 AG077700 (JM) and the Undergraduate Research Opportunities Program (UROP) at University of California, Irvine. We thank Jeffrey Kim, Pathology and Laboratory Medicine, University of California-Irvine for immunostaining of brain sections. We acknowledge the use of QuPath for digital analysis of IHC.

